# MMETHANE: interpretable AI for predicting host status from microbial composition and metabolomics data

**DOI:** 10.1101/2024.12.13.628441

**Authors:** Jennifer J. Dawkins, Georg K. Gerber

## Abstract

Metabolite production, consumption, and exchange are intimately involved with host health and disease, as well as being key drivers of host-microbiome interactions. Despite the increasing prevalence of datasets that jointly measure microbiome composition and metabolites, computational tools for linking these data to the status of the host remain limited. To address these limitations, we developed MMETHANE, an open-source software package that implements a purpose-built deep learning model for predicting host status from paired microbial sequencing and metabolomic data. MMETHANE incorporates prior biological knowledge, including phylogenetic and chemical relationships, and is intrinsically interpretable, outputting an English-language set of rules that explains its decisions. Using a compendium of six datasets with paired microbial composition and metabolomics measurements, we showed that MMETHANE always performed at least on par with existing methods, including blackbox machine learning techniques, and outperformed other methods on >80% of the datasets evaluated. We additionally demonstrated through two cases studies analyzing inflammatory bowel disease gut microbiome datasets that MMETHANE uncovers biologically meaningful links between microbes, metabolites, and disease status.

## INTRODUCTION

The human gut microbiota is an extremely complex ecosystem that performs a range of critical functions for the host [1-3]. Alterations affecting functioning of the microbiome have been associated with many human diseases and disorders including, allergies, inflammatory bowel disease (IBD), cardiovascular disease, chronic kidney disease, metabolic diseases such as diabetes, and infections [3-5]. Although many such associations have been reported, understanding of the specific molecular mechanisms through which the microbiome influences host health or disease remains limited.

Metabolites represent one very important mechanism for microbe-microbe and host-microbiome crosstalk. In the gastrointestinal tract in particular, metabolic activity is intense, with production, absorption, and modification of a myriad of compounds by both microbial and host cells [1, 6]. Indeed, strong mechanistic links have been established between some human diseases and microbially derived metabolites, such as promotion of cardiovascular disease by trimethylamine N-oxide [7]; protection against intestinal inflammation in IBD and *Clostridioides difficile* infection (CDI), by butyrate [8]; and protection against CDI by amino acid depletion [9, 10]. These well-studied metabolites likely represent the tip-of-the-iceberg, with high-throughput metabolomics and metagenomics methods offering promise as platforms for discovery of novel metabolite-mediated relationships between the microbiome and host health and disease. Such data, which is high dimensional, noisy, and often has low samples sizes, requires rigorous computational analyses [11]. Moreover, to be most useful for discovering and ultimately understanding underlying mechanisms of host-microbial interactions, it is import that computational analysis methods produce human-interpretable results [12].

Existing computational methods for discovering relationships between high-throughput metabolomics/metagenomics data and host status have several shortcomings. One of the most popular methods is univariate hypothesis testing [4, 13-16], which evaluates differences between cases and controls for each microbial or metabolic feature of interest. These analyses, although straightforward to implement and interpret, consider each feature individually, and thus cannot identify relationships that arise from interactions among multiple features [17], and also have limited statistical power due to the need for multiple hypothesis correction [18]. Supervised machine learning methods are an alternative, which unlike univariate hypothesis testing approaches, can consider all features simultaneously and predict unseen data [17]. Examples of supervised machine learning methods that have been used to analyze metabolomics and metagenomic data include lasso logistic regression (LR) and random forests (RFs) [5, 19, 20]. Popular supervised machine learning methods that have been used extensively in other fields include boosting and deep neural networks (DNNs). An important trade-off for many supervised machine learning methods is their ability to capture complex, nonlinear relationships that may occur in biological systems versus the degree to which their decisions are human-interpretable. For example, the nonlinear methods RFs, boosting, and DNNs, are not directly interpretable, requiring post hoc analyses to understand model choices. These analyses are not faithful to the underlying models and offer explanations that may be oversimplifications or distortions of how the model maps input features to predictions [21].

To address the above-mentioned challenges for linking metabolomic and metagenomic measurements to host status, we developed a computational method, MMETHANE (Microbes and METabolites to Host Analysis Engine), which we make available as an open-source software package. MMETHANE builds on our prior supervised machine learning methods, MITRE and MDITRE [22, 23], which learn human-interpretable rules for predicting a binary label (host status) from longitudinal microbial sequencing data (16S rRNA amplicon or shotgun metagenomics) and the phylogenetic relationships among the taxa. The key advance over our previous methods is that MMETHANE also analyzes metabolomic data and does so *jointly* with microbial composition data. To accomplish this, we modeled inter-metabolite relationships using structure-based chemical distance measures. This domain-specific knowledge enables the model to perform adaptive, biologically relevant dimensionality reduction, yielding interpretable predictors that can also potentially increase statistical power in the setting of noisy data and low sample sizes.

The remainder of this manuscript is organized as follows. We first present the MMETHANE method, describing the purpose-built model and its incorporation of domain-specific knowledge. Next, we detail the compendium of six datasets we compiled to benchmark MMETHANE and comparator methods. Using these datasets, we assess five structure-based chemical similarities for MMETHANE, to determine which is most informative for use in subsequent benchmarking. We then compare predictive performance of our method against four other supervised learning methods—LR, RFs, AdaBoost, and DNNs—on the dataset compendium. To further understand observed differences in predictive performance between the methods, we benchmark them on semi-synthetic data with varying sample sizes and types of associations that may occur in real data. Finally, we assess MMETHANE’s ability to uncover biologically meaningful relationships among metabolites, microbes, and the host, through two case studies.

## RESULTS

### MMETHANE is a purpose-built deep learning method provided as an open-source software package that infers human-interpretable rules predicting host status from microbial composition and metabolomics data

MMETHANE is a supervised machine learning method that predicts host status (e.g., disease or no disease) from paired microbial composition and metabolomic data (**Figure 1A**). The MMETHANE model captures non-linear interactions while maintaining human-interpretability through a custom architecture that learns sparse sets of English-language *rules*. As shown in **Figure 1B**, MMETHANE uses a feedforward deep neural network with four layers. The parameters learned by the layers can be directly interpreted as components of rules. Each rule consists of a set of *detectors*. Each detector learns: (a) a group of microbial taxa or metabolites and aggregation operation for the group, i.e., sum of relative abundances of taxa or averages of standardized metabolite levels (layer *i*), and (b) an activation threshold, i.e., the detector is *on* if the aggregated value for the group surpasses the threshold (layer *ii*). Activations for the detectors are combined via a logical-AND to form the activation for each rule (layer *iii*). Finally, a prediction for the status of each host is calculated from a weighted combination of rule activations (layer *iv*). MMETHANE uses a Bayesian framework to bias the model toward sparsity, i.e., learning small numbers of detectors and rules. Because efficiently learning parameters in rule-based models is generally very computationally intensive [23], we developed an efficient inference method based on mathematical relaxations of discrete-valued functions. See **Methods** for details.

**Figure 1.**
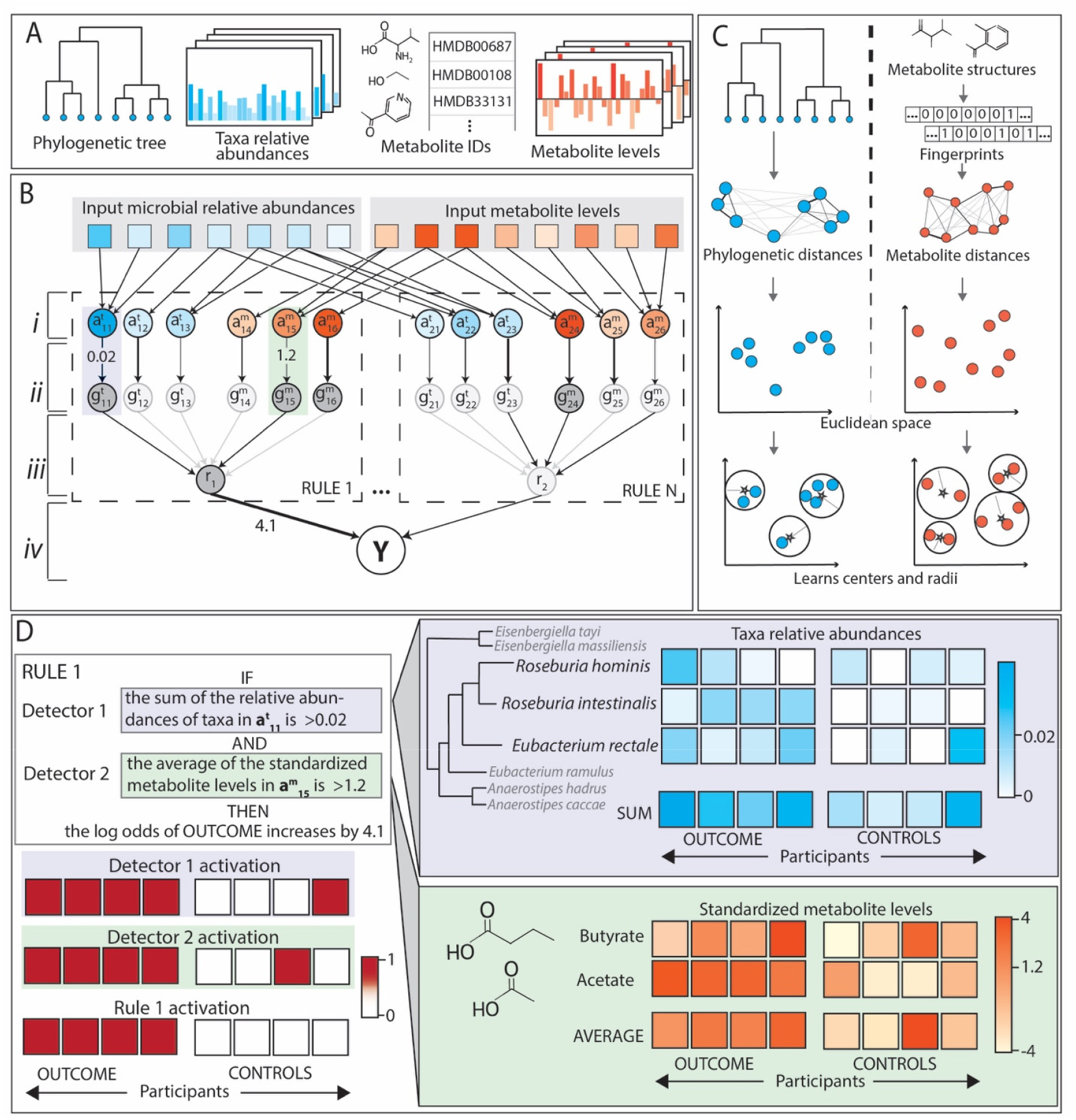
MMETHANE is a purpose-built deep learning method that infers human-interpretable rules predicting host status from microbial composition and metabolomics data. **(A)** Inputs to MMETHANE are a phylogenetic tree, microbial relative abundances, metabolite IDs (from which structures are automatically retrieved), and metabolite levels. **(B)** Schematic of the custom deep learning architecture. Input taxa and metabolites are processed by a feedforward neural network with four layers: *(i)* feature focus, aggregates input features (taxa or metabolites) into detectors, *(ii)* detector activation, “on” (dark grey) if the input from the previous layer surpasses a threshold, *(iii)* logical-AND layer, combines detector activations into a final rule activation, and *(iv)* prediction layer, outputs the probability of the host status label based on the activations and strengths of the input rules. The model probabilistically selects which detectors and rules to include, with bias toward sparsity (depicted here by the darkness of edges). **(C)** Schematic of how feature focus layers use phylogenetic or metabolite distances to learn groupings for detectors. Distances, derived either from sequences for taxa or chemical structures for metabolites, are embedded into Euclidean spaces, and each detector learns a center and radius that defines the grouping it operates on. **(D)** Example of how the learned model structure is directly interpretable as English-language rules. Edges from the inputs → layer (*i*) define which taxa or metabolites participate in each detector; edges from layer (*i*) → (*ii*) define the detector threshold; edges from layer (*ii*) → (*iii*) define which detectors are used; edges from layer (*iii*) → (*iv*) define which rules are used, and the edge weights define the contribution of the rule to the log odds of the host status label.

As with our previous work on rule-based models for microbiome data analysis [22, 23], we designed the MMETHANE model to leverage prior knowledge to learn biologically informed groupings (**Figure 1C**) of input features (taxa or metabolites). The matrix of pairwise distances between input features are embedded in a Euclidean space, and then each detector learns a center and radius in this space. Inclusion in the group is then determined using a relaxed set inclusion distance-based function that we developed [23]. Distances for taxa are based on phylogenetic trees, as in our previous work [22, 23]. For metabolites, we developed a framework for modeling distances based on molecular fingerprinting approaches. We investigated a range of fingerprinting approaches and include the capability in the MMETHANE software to calculate five different fingerprints from input metabolite IDs. See **Methods** and the subsequent section on empirical choice of an optimal fingerprinting approach for additional details.

We provide MMETHANE to the community as an open-source Python package. To run an analysis using the software, the user supplies four files as inputs: (1) a phylogenetic tree (or a default tree may be used), (2) a table of microbial counts or relative abundances, (3) a table of metabolite IDs, and (4) a table of continuous-valued metabolite levels. See **Methods** and the software documentation for a complete description of the input formats. The outputs of the MMETHANE software are: (1) a binary prediction for each input subject, (2) a set of English-language rules explaining model decisions, and (3) visualizations for each rule. Examples of rules and their visualizations are shown in **Figure 1D** and in the subsequent case-study sections.

### Compendium of paired microbial composition and metabolomics data

To facilitate benchmarking of MMETHANE and state-of-the-art methods, we compiled a compendium of publicly available datasets containing paired microbial sequencing (either 16S rRNA amplicon or shotgun metagenomics) and metabolomics measurements. Most datasets included in the compendium were part of the curated gut microbiome-metabolome data resource [24]. To ensure bioinformatics consistency, microbial sequencing data was reprocessed using dada2 for 16s rRNA amplicon data [25], and Metaphlan3 for metagenomics data [26]. We limited datasets chosen to those with at least two distinct outcome/treatment groups, as our goal was to assess predictive accuracy of methods. After running benchmarking (see below), we discarded for further consideration any datasets in which no method achieved an area under the receiver-operator curve (AUC) > 0.6 on either metabolomics or microbial sequencing data (or combined), as those datasets were deemed to have too little signal to be used to compare performance meaningfully across methods.

This selection procedure yielded six datasets for the compendium: He et al. [13], Dawkins et al. [5], Erawijantari et al. [16], Lloyd-Price et al. [14], Franzosa et al. [15], and, Wang et al. [4]. Briefly, He et al analyzed fecal samples from breastfed and formula-fed infants using H1 NMR metabolomics and 16S rRNA amplicon sequencing [13]. Dawkins et al analyzed fecal samples of patients diagnosed with *Clostridioides difficile* infection (CDI) after completion of antibiotic treatment using 16S rRNA amplicon sequencing and LC-MS metabolomics [5]. Erawijantari et al analyzed fecal samples from 42 participants who had previously undergone gastrectomy and 54 healthy controls using shotgun metagenomics and time-of-flight MS (TOF-MS) [16]. Lloyd-Price et al analyzed fecal samples with LC-MS metabolomics and shotgun metagenomics from 79 IBD patients and 26 healthy controls [14]. Franzosa et al. used the same methodology as Lloyd-Price et al to analyze fecal samples from 121 IBD patients and 34 healthy controls [15]. Wang et al performed HPLC-MS/MS metabolomics and shotgun metagenomic sequencing on fecal samples from 220 patients with end-stage renal disease (ESRD) and 67 healthy controls [4]. Additional details on each dataset can be found in **Supplementary Table 1**.

### Benchmarking on the data compendium selected the PubChem CACTVS fingerprint as the best metabolite measure for MMETHANE

To assess the influence of the molecular fingerprinting measures on predictive performance in MMETHANE, we evaluated five fingerprints: PubChem CACTVS [27], Morgan, MAP4 [28], MQN [29] and InfoMax [30]. We chose these fingerprints based on their prevalence of usage in the literature and to span a range of complexities and approaches, including molecular fingerprints based on substructure features, metabolite topology, and deep learning representation of metabolite features. See **Methods** and **Supplementary Table 2** for additional details on the fingerprints we evaluated. Similarities were calculated from fingerprints using the Tanimoto/Jaccard metric for the binary PubChem CACTVS and Morgan fingerprints, and the city block distance for the MQN fingerprint; for the MAP4 and InfoMax fingerprints, we used the similarities provided by their respective software packages. For each fingerprinting method, we evaluated MMETHANE’s 5-fold cross-validated predictive performance (AUC) on the data compendium.

Overall, the PubChem CACTVS fingerprint provided the most consistent performance (**Supplementary Figure 1**). It was the top performer in 3/6 datasets: Dawkins et al., Erawijantari et al., and Fransoza et al. For the other three datasets, although there were a few instances in which performance with the PubChem CACTVS fingerprint was not significantly different than with other fingerprints, no other fingerprint outperformed it with statistical significance. Thus, for efficiency and compactness of our presentation, we employ the PubChem CACTVS fingerprint for the subsequent analyses described. However, as described above, we provide the option in the MMETHANE software for users to easily calculate similarities with any of the other methods, to assess performance on their own data.

### MMETHANE predicted host status as well or better than state-of-the-art machine learning methods on the data compendium

We benchmarked MMETHANE’s predictive performance on all datasets in the compendium against that of four popular supervised machine learning methods: lasso logistic regression (LR), Random Forests (RF), AdaBoost, and a feed-forward neural network (FFNN). LR was chosen as a comparator method because it has been widely used for analyzing microbiome data and is inherently interpretable [5, 19, 20]. RF, a nonlinear method, has also been widely used for analyzing microbiome data, including on datasets with paired microbial sequencing and metabolomics measurements [5, 19, 20]. AdaBoost and FFNN are exemplars of additional types of nonlinear methods, and are very popular in the broader machine learning field, with some applications in the microbiome field, as well [31-37]. Note that the nonlinear methods RF, AdaBoost, and FFNN are not inherently interpretable, and post hoc approaches for their interpretation are not generally faithful to the underlying models, which can result in incorrect inferences [38]. We evaluated performance using five-fold cross-validated AUC, as described above, and on three different prediction tasks: metabolites only, taxa only, and metabolites and taxa combined.

Overall, MMETHANE performed on par or outperformed all the comparator methods (**Figure 2**), across all prediction tasks. For predicting from metabolomics data only, MMETHANE was the top performer on 4/6 datasets. On microbial composition data only, MMETHANE and FFNN were top performers pn 3/4 datasets (note that 2 datasets were excluded from this analysis, because no method achieved an AUC above 0.6). On combined metabolomic and microbial composition data, MMETHANE was the top performer on 5/6 datasets. Notably, MMETHANE outperformed the only other directly interpretable method, lasso LR, in all cases except for predicting CDI recurrence from metabolomics data only.

**Figure 2.**
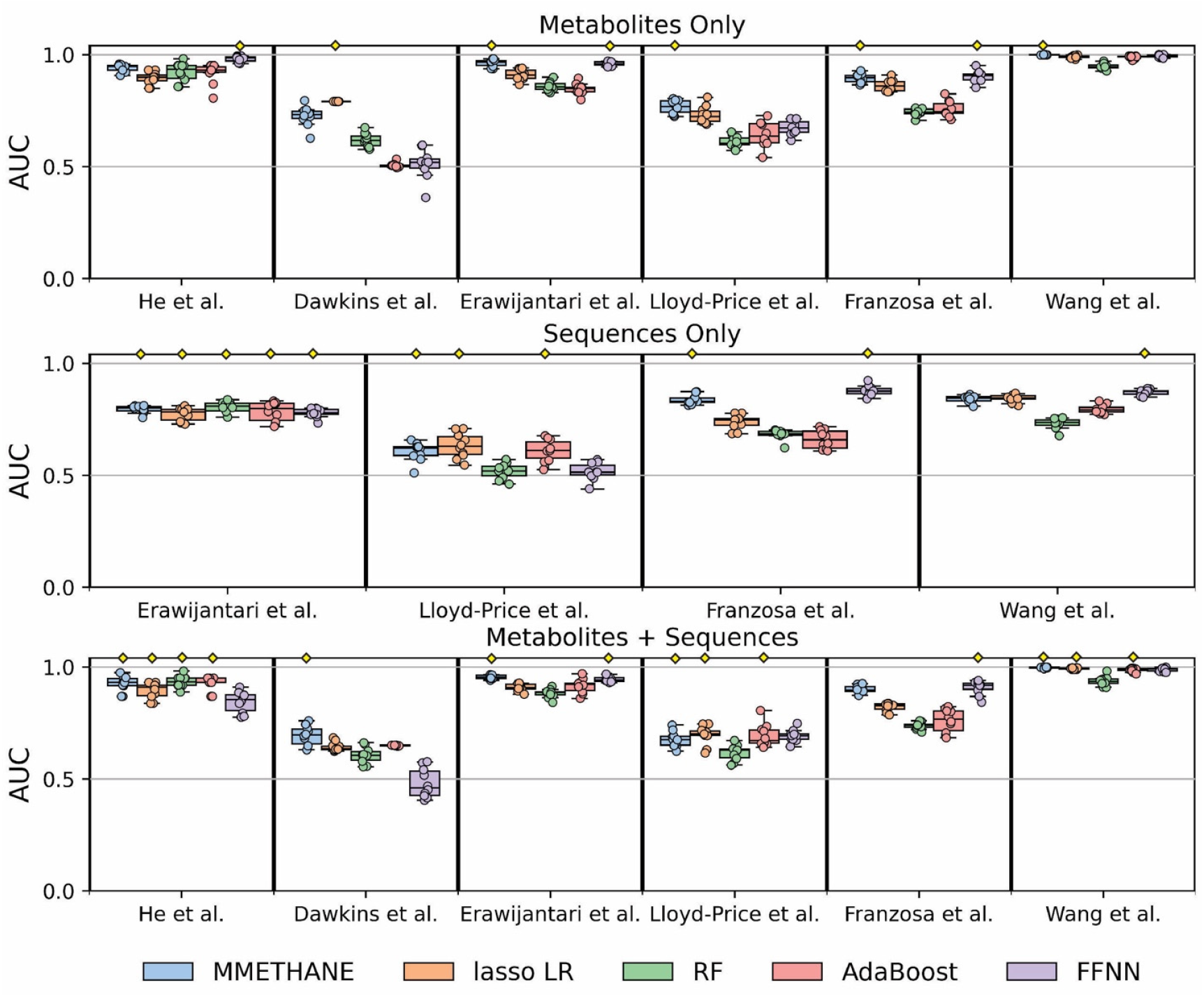
MMETHANE predicted host status on par or better than all state-of-the-art comparator methods on the compendium of microbial sequencing and metabolomics data. Five-fold cross-validated AUC scores for prediction of host status on the six datasets in the compendium are shown, for MMETHANE, lasso logistic regression (LR), random forest (RF), adaptive boosting (AdaBoost), and a feed forward neural network (FFNN). Methods were evaluated with inputs of metabolomic data alone, sequence data alone, or both. Box plots with results from ten random seeds, with medians and 95% intervals, are shown. Yellow diamonds indicate the top score or scores (if multiple scores were not significantly different from the top score).

Our benchmarking results also demonstrate that metabolites were more predictive of host status than microbial taxa compositions in these datasets. Moreover, in almost all cases, predictive performance when both inputs were provided was not significantly better than performance when metabolites only were provided. This result is consistent with prior observations regarding the predictiveness of the gut metabolome versus microbial composition [5]. Interestingly, however, on 5/6 of the datasets, at least one method used both metabolite and microbial composition predictors, and on half the datasets, all methods used both types of predictors (**Supplementary Table 3**). These results suggest that microbial composition data, although not evidently synergistic with metabolomics data for predicting host status, can still provide relevant signal.

### MMETHANE exhibited consistently superior performance across sample sizes and types of associations compared to state-of-the-art methods as assessed on semi-synthetic data

We next sought to understand the basis for MMETHANE’s strong performance on the data compendium. Factors that we reasoned could be responsible for this included MMETANE’s ability to detect combinations of biologically plausible associations, and data efficiency (ability to perform well with small sample sizes while continuing to increase performance with more data). An analysis of these factors requires ground truth knowledge of associations, so we employed a semi-synthetic data generation approach that artificially introduced differences into data from the control group of a reference dataset. For the reference dataset, we used Wang et al. [4], because it had the largest number of control samples in the data compendium.

Briefly, our semi-synthetic dataset generation and evaluation procedure was as follows. For each simulation, half the participants were selected (uniformly at random, with replacement) to be “cases” that received the perturbation and half were selected (uniformly at random, with replacement) as “controls” that did not. Next, a group (or groups) of metabolites and/or microbial taxa were selected uniformly at random to be perturbed using parameters derived from the original data (e.g., effect sizes of perturbations). We considered five different perturbation scenarios that assessed methods’ abilities to predict from metabolomic or taxa data separately, or in combination (**Figure 3**). The first and third scenarios, which involved perturbing either a single metabolite group or a single taxonomic clade, were expected to be the easiest cases for the algorithms to learn. The remaining scenarios included combinations of associations (e.g., as shown in **Figure 3A**, for associations with both microbial compositions and metabolites), emulating biological phenomena that involve nonlinear interactions such as allostery [39]. These types of relationships were expected to be more difficult for the algorithms to learn; in particular, logistic regression, which does not model interactions among variables, was expected to perform poorly on these cases. For all the scenarios, we assessed predictive performance on datasets of varying sample sizes. We simulated data with 36, 48, 64, 128, 300, and 1000 participants; this range was selected based on realistic dataset sizes with the lower and upper values smaller or larger than those of the datasets analyzed. We evaluated MMETHANE against the four comparator methods and using the same cross-validation strategy as for the real data as described above. See **Methods** for full details on the semi-synthetic data generation procedure.

**Figure 3.**
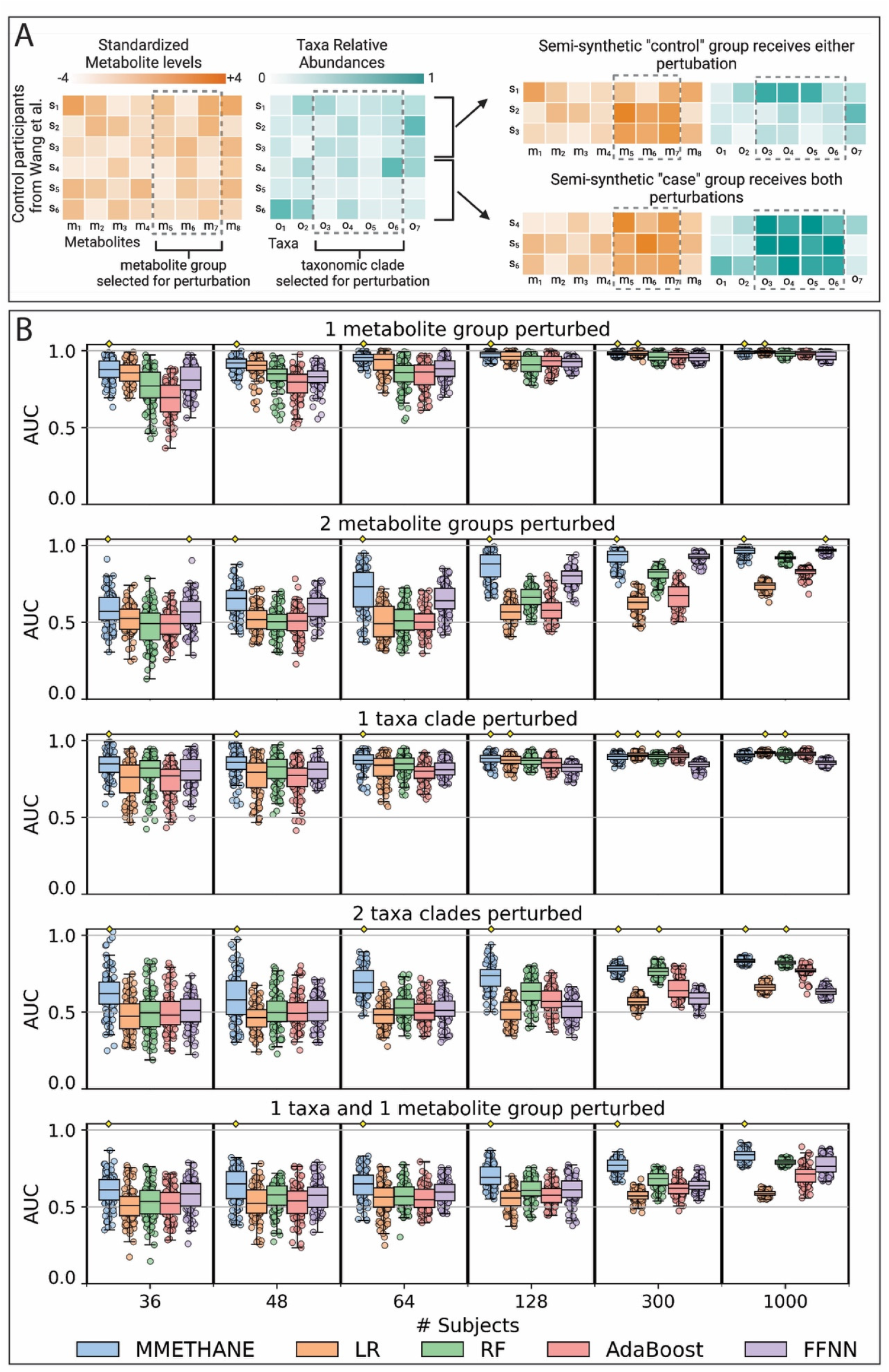
MMETHANE exhibited consistently superior performance across sample sizes and types of associations compared to state-of-the-art methods assessed on semi-synthetic data. A semi-synthetic data generation approach was used that introduced simulated differences into a real reference dataset. Five scenarios were created to assess methods’ abilities to predict from metabolomic or taxa data separately, or in combination, and in each scenario the sample size was varied to assess data efficiency of the methods. **(A)** Schematic showing the simulation process for scenario 5, the most complex, in which 1 metabolite and 1 taxa group were perturbed. Participants from the control group of a reference real dataset were randomly assigned to synthetic “control” and “case” groups. Members of the “control” group then received either the metabolite group or the taxa group perturbation, but not both, whereas members of the “case” group received both types of perturbations. Scenarios 2 and 4 were generated similarly, except they involved 2 groups of either metabolites or taxa, but not both. Scenarios 1 and 3 involved a single group of either metabolites or taxa. (**B)** Five-fold cross-validated AUC scores for prediction of host status for the five scenarios are shown, for MMETHANE, lasso logistic regression (LR), random forest (RF), adaptive boosting (AdaBoost), and a feed forward neural network (FFNN). Ten random semi-synthetic datasets were generated for each scenario and sample size, and models were run with ten random seeds. Box plots indicate medians and 95% intervals. Yellow diamonds indicate the top score or scores (if multiple scores were not significantly different from the top score).

MMETHANE outperformed or was on par with all the other methods on all the semi-synthetic data scenarios (**Figure 3B**). Some general trends emerged, which were useful in understanding performance differences among the methods. First, comparing across scenarios with metabolomic data, taxa composition data, or both, we saw that all methods generally performed better on the metabolite data alone. This was consistent with our results on real data and indicated that our semi-synthetic data generation procedure captured this phenomenon. Second, even on the simplest scenarios, of perturbation of a single group of metabolites or a single taxonomic clade, MMETHANE remained the sole top performer up to a sample size of 128 subjects, after which it shared the top spot with multiple methods. These results indicate that MMETHANE was more data efficient than the other methods, likely because it not only encodes penalties on model complexity, but also incorporates biological knowledge through known phylogenetic or chemical relationships. Third and finally, as expected, all the methods, including MMETHANE, made less accurate predictions on the more complex scenarios that involved interactions between sets of taxa, metabolites, or both. Also, as expected, LR performed poorly on these scenarios, even with large sample sizes, because it is incapable of capturing nonlinear interactions. Interestingly, methods capable of capturing nonlinear interactions other than MMETHANE did not show consistent improvements in performance with larger sample sizes, as they had on the simpler scenarios.

### Case studies: MMETHANE learned rules linking gut bacterial taxa and metabolites to inflammatory bowel disease in the host

After demonstrating that MMETHANE performed on par or outperformed other supervised machine learning methods on real and semi-synthetic data, we next investigated our method’s ability to discover biologically meaningful associations. We used data from two analyses of gut microbes and metabolites from participants with IBD as our case studies. Our rationale for choosing these studies was the clinical importance of IBD and its complexity as a pathological entity, with the rationale that MMETHANE’s ability to group taxa or metabolites and find interpretable non-linear relationships between them could provide additional insights into underlying mechanisms of microbe, metabolite, and host interactions in the disease.

#### Case Study 1: Lloyd-Price et al

MMETHANE found two rules on this dataset (**Figure 4A-B**), which analyzed fecal samples with LC-MS metabolomics and shotgun metagenomics from 79 participants with IBD and 26 healthy controls. The first rule, which contains a single detector, states that if the aggregated relative abundance of a group of 17 taxa is increased above the learned threshold, the odds of the sample being classified as IBD is decreased by 3 × 10^5^ fold (**Figure 4A**). Many of the taxa have previously been associated with anti-inflammatory activity and intestinal barrier integrity [7, 40], and are producers of short chain fatty acids, which are known to mediate these effects [8, 41, 42]. This rule thus identifies taxa with a biologically plausible protective role in IBD, consistent with its semantics that higher levels of these taxa are associated with lower odds of IBD.

**Figure 4.**
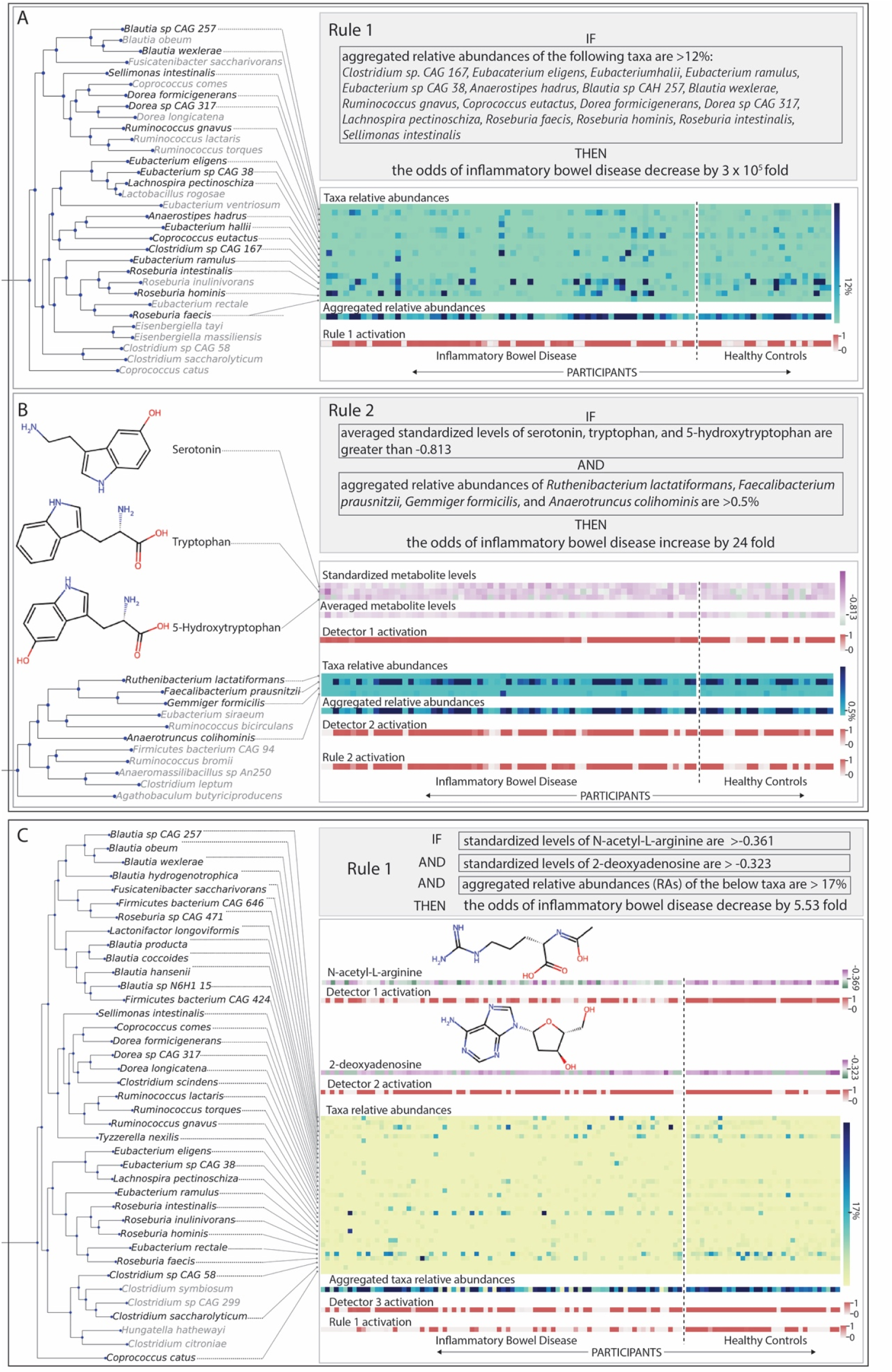
MMETHANE learned interpretable rules with biologically relevant taxa and metabolites for classifying inflammatory bowel disease (IBD) versus healthy controls. Visualizations automatically produced by the software are shown. Each visualization includes the phylogenetic relationships of the identified taxa and the chemical structures of the identified metabolites, as well as a display of the underlying data and how the rule and detector activations influence the prediction for each participant. **(A)** and **(B)** show the two rules MMETHANE learned when applied to the Lloyd-Price et al. dataset, which analyzed fecal samples with LC-MS metabolomics and shotgun metagenomics from 79 participants with IBD and 26 healthy controls. The first rule **(A)** consists of a single detector (clause), which states that if the aggregated relative abundance of a set of 17 taxa, many of which are short-chain fatty acid producers, is greater than the threshold, then the odds of the participant having IBD is lower. The second rule **(B)** consists of two detectors (clauses), which state that if the aggregated relative abundance of a group of 4 taxa is greater than a threshold AND the average levels of a group of three metabolites is greater than a second threshold, then the odds of the participant having IBD is increased. As discussed in the text, this rule suggests a possible context dependent role for the identified group of bacteria, which may modulate production of pro-inflammatory serotonin. **(C)** Shows the rule MMETHANE learned when applied to the Franzosa et al. dataset, which analyzed fecal samples with LC-MS metabolomics and shotgun metagenomics from 121 participants with IBD and 34 healthy controls. The rule consists of three detectors (or clauses), which if all are true predicts the odds of the participant having IBD is decreased. The first two detectors identify single metabolites, and the third detector identifies a group of 35 taxa. As discussed in the manuscript, the rule identifies metabolites and taxa with possible synergistic roles in promoting a non-inflammatory intestinal environment.

The second rule contained one metabolite detector and one taxa detector, and states that if the aggregated values of both detectors are above the learned thresholds, the odds of the sample being classified as IBD is increased by 24-fold (**Figure 4B**). The metabolite detector in this rule groups together three compounds, tryptophan, 5-hydroxy-tryptophan, and serotonin, which, are the substrate, intermediate, and product of serotonin biosynthesis from tryptophan, respectively. Human and animal studies have shown that increased levels of serotonin resulted in increased severity of intestinal colitis through inhibition of autophagy and beta-defensin production in gut epithelial cells [43, 44], providing a possible mechanism explaining this detector’s utility in predicting IBD. The microbial detector in this rule identified a group of four taxa, *Anaerotruncus colihominis, Faecalibacterium prausnitzii, Gemmiger formicilis*, and *Ruthenibacterium lactatiformans*. Although these taxa are SCFA producers [41, 45, 46], which as discussed for rule one, typically are associated with protection against IBD, rule two suggests these taxa may also play a more nuanced role in IBD in the context of increased serotonin levels in the gut. Interestingly, microbially-derived SCFAs have been found to upregulate Tph1 and thus increase serotonin synthesis in gut epithelial cells [44]. Additionally, tryptophan, the substrate of serotonin biosynthesis is metabolized by bacteria in the gut, including, *Ruminococcus gnavus* and *Eubacterium halii* [47, 48], which were identified in the first rule. These findings thus suggest a possible link between serotonin biosynthesis in the gut and microbial activities that may modulate both substrate availability as well as serotonin production in host epithelial cells and its promotion of inflammation.

#### Case Study 2: Franzosa et al

MMETHANE found a single rule for this dataset (**Figure 4C**), which analyzed fecal samples with LC-MS metabolomics and shotgun metagenomics from 121 participants with IBD and 34 healthy controls. The rule, which contained three detectors (two metabolite and one microbial detector), states that if the aggregated values of all detectors are greater than the learned thresholds, then the odds of the sample being classified as IBD is decreased by 5.5-fold. The first metabolite detector in the rule identified 2-deoxyadenosine, a precursor of dATP (and ultimately DNA), and thus may be a marker of epithelial cell proliferation or commensal bacterial growth. The second metabolite detector in the rule identified N-acetyl-L-arginine, a precursor in arginine biosynthesis, which may similarly be a marker of healthy cellular growth. Additionally, arginine has been shown to increase intestinal barrier integrity and reduce susceptibility to colitis [49-52]. The microbial detector in the rule identified 35 taxa, including species within the *Blautia, Roseburia, Dorea, Ruminococcus*, and *Eubacterium* genera. A number of these taxa overlap with those in the first rule in case study 1, and are SCFA producers, including *Eubacterium eligens, Roseburia intestinalis, Roseburia hominis, Dorea formicigenerans*, and *Sellimonas intestinalis*. Interestingly, there is a possible mechanistic link between this microbial detector and the second metabolite detector: relative abundances of *Ruminococcus* and *Clostridia* species [50] have been shown to be higher with increased levels of arginine, and two taxa in the microbial detector, *Ruminococcus gnavus* and *Clostridium scindens* are known fermenters of alpha-amino acids, including arginine [10, 48, 53]. Taken together, this rule identified metabolites and taxa with probable synergistic roles in promoting a healthy, non-inflammatory intestinal environment.

## DISCUSSION

We have presented MMETHANE, a purpose-built deep learning method that incorporates domain-specific knowledge to learn human-interpretable rules that predict the status of the host from paired metabolomics and microbial composition data. Our results on a compendium of real data demonstrated that MMETHANE’s predictive performance was on par with or surpassed that of state-of-the-art supervised learning methods, including black-box approaches. Analyses on semi-synthetic data moreover demonstrated that MMETHANE was more data efficient than the comparator methods, including on complex scenarios modeling nonlinear relationships among metabolites and/or taxa. Further, we demonstrated that the human-readable rules output by MMETHANE, which automatically group together phylogenetically similar taxa or biochemically-related metabolites, provided biologically relevant insights and suggested interesting hypotheses about the complex interplay between microbes, metabolites, and the host. For example, MMETHANE learned a rule from a dataset with gut metagenomics and metabolomics data from participants with IBD, which suggested a hypothesis that serotonin may have a role in IBD that is modulated by different groups of microbes that either alter substrate availability or have effects on serotonin production in host epithelial cells.

This study has several limitations that suggest future work. First, MMETHANE does not currently incorporate time-series analysis capabilities, as did our previous methods [22, 23] upon which MMETHANE is based. MMETHANE could be readily extended to handle longitudinal datasets, in fact; the reason we did not do so in the present study was due to the lack of available paired metabolomics and microbial composition time-series datasets with host outcomes which would be needed to evaluate our method. Indeed, such datasets would be very valuable for the field overall, as they could provide potentially causal insights into microbe-microbe and host-microbiome interactions, which are difficult to resolve in cross-sectional studies. Second, although not directly related to the MMETHANE model itself, we found that for at least two of the datasets we used as benchmarks, He et al. and Wang et al., the levels of a small number of metabolites (i.e., from specific dietary components, medications, or the disease process itself) almost completely differentiated the participant groups under study. This highlights the need for future studies with larger cohort sizes and designs that allow for explicit control or modeling of confounding factors, which will be important for establishing the relevance of specific gut metabolites to host diseases. Third, MMETHANE is a fully supervised machine learning method, which means that by design it is biased toward finding factors, and interactions among those factors, that influence host status. This approach has the advantage of potentially yielding more accurate predictors, but has the disadvantage of potentially missing biologically relevant interactions that do not directly impact predictive accuracy. In future work, it would be interesting to investigate whether adding a generative modeling component to MMETHANE upstream of its prediction layers (e.g., including latent variables that jointly generate the taxa composition and metabolomic data) could effectively balance predictive accuracy with modeling other types of relationships in the data. Fourth and finally, the detector structure of MMETHANE that incorporates biological knowledge, although demonstrably effective on the datasets analyzed, could be extended to capture richer information. For example, MMETHANE currently uses phylogenetic trees constructed from 16S rRNA gene sequences. Future models could incorporate the full gene content of organisms, which could help the method to group together phylogenetically distant but functionally related organisms. Additionally, such information could be layered with knowledge of metabolic pathways, which could allow the model to explicitly link microbes and their metabolic inputs and outputs.

## CONCLUSION

MMETHANE, which we make available as an open-source software package, is a computational tool for discovering predictive relationships between paired high-throughput microbial sequencing and metabolomics data, and the status of the host. Our method employs a custom-designed deep learning architecture incorporating biological knowledge to provide accurate predictions without sacrificing interpretability. We thus believe MMETHANE will be a valuable resource for researchers seeking to elucidate the interplay between microbes, metabolites, and the host.

## ACKNOWLEDGEMENTS

We would like to thank Liat Shenov and Travis Gibson for helpful feedback and discussions about the MMETHANE model and datasets analyzed.

This work was supported by NSF MTM2 2025512, NIH NIGMS R01GM130777, NIH NIGMS R35GM149270, The Massachusetts Life Sciences Center, and the Brigham and Women’s Hospital President’s Scholar Award.

## AUTHOR CONTRIBUTIONS

Conceptualization, G.K.G. and J.D.; Methodology, G.K.G. and J.D.; Software, J.D.; Validation, J.D.; Formal Analysis, J.D.; Investigation, J.D.; Resources, G.K.G; Writing, G.K.G. and J.D.; Visualization, J.D..; Supervision, G.K.G.; Funding Acquisition, G.K.G.

## METHODS

### KEY RESOURCE TABLE

| REAGENT or RESOURCE | SOURCE | IDENTIFIER |
| --- | --- | --- |
| **Deposited data** |  |  |
| 16S rRNA sequencing samples from He et al. [13] | European Nucleotide Archive (ENA) | ERP112481 |
| Metabolomics data from He et al. [13] | The curated gut microbiome-metabolome data resource (https://github.com/borenstein-lab/microbiome-metabolome-curated-data.wiki.git) [24] | HE-INFANTS-MFGM-2019 |
| 16S rRNA sequencing samples from Dawkins et al. [5] | Sequence Read Archive (SRA) | PRJNA772946 |
| Metabolomics data from Dawkins et al. [5] | [5] | [10.5281/zenodo.6385654](https://doi.org/10.5281/zenodo.6385654) |
| Metagenomics sequencing data from Erawijantari et al. [16] | DNA Data Bank of Japan (DDBJ) Sequence Read Archive (DRA) | DRA007281, DRA008243, DRA006684 and DRA008156 |
| Metabolomics data from Erawijantari et al. [16] | The curated gut microbiome-metabolome data resource (https://github.com/borenstein-lab/microbiome-metabolome-curated-data.wiki.git) [24] | ERAWIJANTARI-GASTRIC-CANCER-2020 |
| Metagenomics sequencing data from Lloyd-Price et al. [14] | Sequence Read Archive (SRA) | PRJNA398089 |
| Metabolomics data from Lloyd-Price et al. [14] | Metabolomics Workbench (http://www.metabolomicsworkbench.org) | PR000639 |
| Metagenomics sequencing data from Franzosa et al. [15] | Sequence Read Archive (SRA) | PRJNA400072 |
| Metabolomics data from Franzosa et al. [15] | Metabolomics Workbench (http://www.metabolomicsworkbench.org) | PR000677 |
| Metagenomics sequencing data from Wang et al. [4] | European Bioinformatic Institute (EBI) database | PRJNA449784 |
| Metabolomics data from Wang et al. [4] | MetaboLights database (http://www.ebi.ac.uk/metabolights/) | MTBLS700 |
| Data to reproduce figures | This paper | [10.5281/zenodo.14064091](https://doi.org/10.5281/zenodo.14064091) |
| Results of machine learning models run | This paper | 10.5281/zenodo.14064001 |
| **Software and algorithms** |  |  |
| MMETHANE model | https://github.com/gerberlab/mmethane [54] | N/A |
| Code to reproduce paper figures | https://github.com/gerberlab/mmethane-paper-results [55] | N/A |
| Metaphlan v3.0 | [56], [26] | N/A |
| DADA2 v1.20 | [25] | N/A |

### RESOURCE AVAILABILITY

#### Lead contact

Further information and requests for resources and reagents should be directed to and will be fulfilled by the lead contact, Georg K. Gerber.

#### Materials availability

This study did not generate any unique materials or reagents.

#### Data & Code availability

The MMETHANE model is available at https://github.com/gerberlab/mmethane [54]. Code to reproduce all analyses and figures in this project, as well as the semi-synthetic and real datasets used to produce results is available at https://github.com/gerberlab/mmethane-paper-results [55]. Data needed to reproduce all figures is available at 10.5281/zenodo.14064091.

## METHOD DETAILS

### MMETHANE software package

MMETHANE is available as an open-source Python software package that reads in input data, trains the model, and provides tools for visualization and interpretation of outputs.

Required and optional inputs and their file formats are as follows:

(1) **Phylogenic tree *(optional, if default is used)*:** a phylogenetic tree newick file (.nwk) or a table of pairwise phylogenetic distances between taxa (.csv, .tsv).
(2) **Metabolite identification numbers** OR **matrix of metabolite similarities/distances *(required)*:** the software will automatically compute metabolite similarities (see below) given a (.csv, .tsv)-formatted table of metabolites and IDs for each metabolite measured. Identification numbers in any of the following formats are accepted: Human Metabolite DataBase (HMDB), KEGG, InChI, InChI Key, or PubChem. Alternatively, the user can provide a table (.csv, .tsv) of pre-computed pairwise distances or similarities between metabolites.
(3) **Microbial counts/relative abundances *(required)*:** a (.csv, .tsv)-formatted table, or a table provided by Metaphlan [26] or Dada2 [25].
(4) **Metabolic measurements *(required)***: a (.csv, .tsv)-table of positive-valued metabolite levels.

MMETHANE supports automatic calculation of metabolite distances using five different molecular fingerprint methods:

(1) **PubChem CACTVS (default)**: a fingerprint of 881-bits, in which each bit corresponds to the presence or absence of a chemical substructure. MMETHANE obtains these fingerprints using the PubChemPy API (v1.0.4) [27].
(2) **Morgan**: also referred to as extended-connectivity fingerprints (ECFP), this fingerprint is calculated by applying a variation of the Morgan algorithm, as described in [57]. MMETHANE obtains these fingerprints from RDKit (v2023.09.2) [58] using rdkit.Chem.AllChem.GetMorganFingerprintAsBitVect with bit-size set to the default of 2048 bits.
(3) **Molecular Quantum Number (MQN)** [29]: a vector of counts for 42 structural features falling into four categories: atoms, bond types, polarity, and topology. MMETHANE obtains these fingerprints using the function rdkit.Chem.rdMolDescriptiors.
(4) **MinHashed atom-pair (MAP4)** [59]: a 1024-bit fingerprint designed to be suitable for small and large molecules by combining elements of atom-pair and substructure fingerprints. MMETHANE obtains these fingerprints using map4 from the GitHub repository [28] with the default radius of 2 and default of 1024 dimensions.
(5) **Infomax** [30, 60]: a 300-bit fingerprint calculated using a pre-trained deep learning model. MMETHANE obtains the fingerprint and distances between the fingerprints by running the pre-trained ‘gin_supervised_infomax’ model [61] (dgllife v0.3.2, dgl v1.1.3) on bigraphs of all metabolites.

MMETHANE outputs cross-validated predictions for each sample as well as a PDF that contains the following:

(1) An English-language description of each rule.
(2) Heatmap visualizations of data used for model decisions.
(3) Visualizations of the phylogenetic relationships of the taxa in each detector.
(4) Visualizations of the metabolic structures of metabolites in each detector.

### MMETHANE model

#### Overview

MMETHANE extends the MITRE/MDITRE models that we previously developed [22, 23], by adding detectors for metabolites. MMETHANE can thus integrate both metabolomic and microbial sequencing data, by learning predictive rules that contain both metabolite and taxa detectors. Note that MMETHANE does not implement the time-series analysis capabilities of MITRE/MDITRE, because sufficient datasets with paired longitudinal metabolite and microbial composition measurements were not available to validate this capability.

The MMETHANE model consists of a purpose-built, Bayesian, feed forward deep neural network (DNN), with the following layers: (i) feature aggregation, (ii) detector activation, (iii) detector selection and logical-AND rule activation, and (vi) rule selection and weighted-OR host status prediction (**Figure 1B**). Prior biological knowledge is incorporated into the model using phylogenetic distances for microbial sequences and distances computed from molecular fingerprints for metabolites. Given the phylogenetic and metabolite distances, and training data consisting of paired metabolite and microbial sequencing data, the DNN learns a sparse set of detectors and rules that are then used to make predictions about the host label.

#### Incorporating phylogenetic distances

MMETHANE uses the same procedure for phylogenetic distance embedding as MDITRE [23]. Briefly, a symmetric *N*_*T*_ × *N*_*T*_ matrix of phylogenetic distances for *N*_*T*_ input taxa is calculated from a reference phylogenetic tree. Embedding into Euclidean space is then performed using Principal Coordinate Analysis (PCoA), with the embedding dimensions *D*_*T*_ automatically selected as the lowest dimension that does not result in a significant difference (*p*<0.05, Kolmogorov-Smirnov (KS) test, SciPy v1.11.3) between the distributions of pre-and post-embedded distances. This yields an embedding matrix, 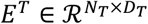 for the taxa.

#### Incorporating metabolite distances

A symmetric *N*_*M*_ × *N*_*M*_ matrix of metabolite distances for *N*_*M*_ metabolites is either provided by the user or calculated by the MMETHANE software package from provided metabolite identification numbers as described above. Distances are then embedded into Euclidean space using PCoA. We did not find that the KS-test yielded practically useful reductions in the dimensionality of the embedding space, as it had for phylogenetic distances. Thus, we chose the embedding dimension 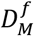 according to the number of components that explained 95% of variance for each fingerprinting method *f*. This procedure yields a set of embedding matrices, 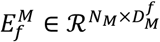.

#### Model layers

To define notation used in the model description below, let *K* denote the total number of possible rules, *L* the total number of possible taxa detectors per rule, and *J* the total number of possible metabolite detectors per rule. As in our previous work, we set *K, J*, and *L* to 10 [22, 23], which we found supports higher than utilized model capacity for the datasets analyzed.

##### (i) Feature aggregation

The first layer learns groups of features (taxa or metabolites) and outputs an aggregate of the features (sum of relative abundances for taxa, mean of averaged standardized levels for metabolites) for each of the learned detectors.

Grouping and aggregation for metabolites was modeled as follows. Let 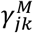 denote the learned center for metabolite detector *j* in rule *k*. As in our previous work [23], we place a Multivariate Normal prior probability distribution with mean 0 and variance 10^4^ on detector centers.

Let 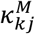 denote the learned radius for detector *j* in rule *k*. For metabolite detector radii, we use a Log-Normal prior calibrated with biological knowledge of metabolites. First, a chemical taxonomy is assigned to each metabolite using Classy Fire [62]. Then, for each metabolite sub-class *c*, we compute the pairwise distance, *q*_*lp*_, between each pair of metabolites in that subclass using the metabolite embeddings *E*^*M*^, and determine the median distance within each sub-class, 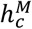 The Log-Normal hyperparameters for the prior distribution on 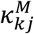 are then computed as the median and variance of the sub-class distances, i.e.:

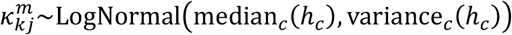

Each detector *j* in rule *k* determines whether to select metabolite *i* according to the detector’s center and radius, using a soft inclusion function:

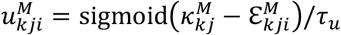

Here, 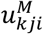 denotes the degree of selection, 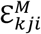 denotes the Euclidean distance in the metabolite embedding space between the detector center and metabolite *i*, and τ_*u*_ is a temperature parameter that controls the sharpness of the selection. As in our previous work [23], we used a linear annealing schedule during training for the temperature parameter from 10^−2^ to 10^−3^.

Finally, for metabolites, the output of the layer is the mean of levels of selected metabolites in each detector:

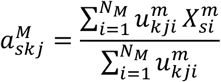

Here, 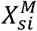 is the log transformed and standardized value of metabolite *i* in subject *s*.

Grouping and aggregation of taxa was modeled for MMETHANE as in our previous work [23]. Briefly, taxa detectors learn centers and radii analogously to the metabolite detectors described. As we did for metabolites, we placed a Multivariate Normal prior on the centers and a Log-Normal prior on the radii. The Log-Normal prior mean and variance hyperparameters were set using median pairwise distances for each taxonomic family. Finally, for taxa, the output of the layer is the sum of relative abundances of selected taxa in each detector:

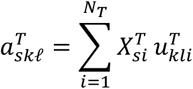

Here, 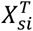 is the relative abundance of taxa *i* in subject *s*.

##### (ii) Detector activation

The second layer takes in the aggregated values for each feature group and learns a threshold of activation for each detector. The layer then outputs a soft binary activation for the detector, depending on whether the feature group aggregate is above or below the learned threshold.

We placed a uniform prior on the metabolite detector thresholds, 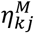, with end-points set based on the values of the metabolites in the data, ***X***^*m*^:

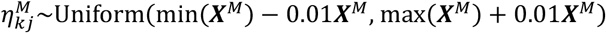

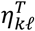, the threshold for each taxa detector, is parameterized by a uniform distribution from 0 to 1, as in [23].

The output of the layer is a soft activation modeled by a sigmoid centered around the learned thresholds for each metabolite or taxa detector:

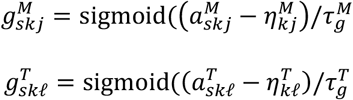

The temperature parameters 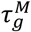 and 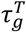 control the sharpness of activation and are annealed during training to ensure final learned values of 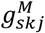 and 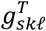 are close to 1 or 0. For taxa detectors, the temperature 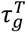 is annealed from 10^−2^ to 10^−3^ during training, as in [23]. The temperature for metabolite detectors, 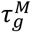, is annealed from 1 to 0.1 during training, because the range of values of 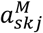 is typically 10 to 100 times larger than the range of values of 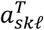.

##### (iii) Detector selection and logical-AND rule activation

The third layer, takes in the outputs of the previous layer, the soft binary activations 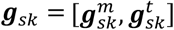 for all possible metabolite and taxa detectors, and selects a sparse set of detectors for the rule. The output of this layer, *r*_*sk*_, is a soft binary activation for each rule *k* and subject *s*, in which the value is approximately 1 when all its selected detectors are approximately 1, as described below.

For each rule k, we denote the set of *J* + *L* detector selectors as 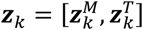. As in our previous work, we place binary concrete priors on the selectors:

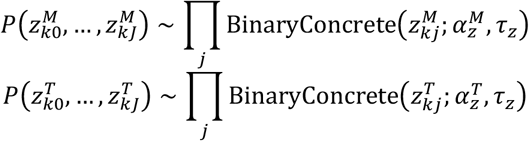

We set the location parameters, 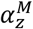 and 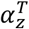, to 1/*J* and 1/*L*, respectively, to encourage sparsity, i.e., a prior assumption of expected values of 1 microbial and 1 metabolite detector per rule. The temperature parameter, τ_*z*_ is annealed from 1 to 0.1 during training, as in [23].

The output of the layer is a soft logical-AND function of the selected detector activations, as in our previous work [23]:

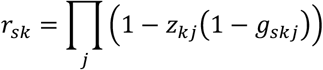

##### (iv) Rule selection and weighted-OR host status prediction

The final layer of the model learns which of the *K* rules to select for the prediction and weights for each selected rule. Analogously to the detector selectors, we placed Binary Concrete priors on the rule selectors ***q***:

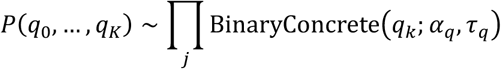

As in our previous work, we set *α*_*q*_ to 1/K to encourage sparsity, i.e., a prior expectation of one rule, and τ_*q*_ was annealed from 1 to 0.1 during training. We placed a diffuse prior on the weights and biases for rules, *β*_*k*_ and *β*_*O*_ (a Normal distribution with mean 0 and variance 1 × 10^*4*^).

The output of the layer for each subject *s* is then the weighted linear combination of selected rule activations and the bias term:

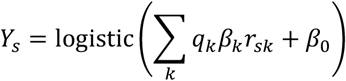

### Inference

We employed maximum a posteriori (MAP) inference to learn the model parameters, using the Adam optimizer in PyTorch (v2.2.0). Initialization of the feature aggregation layer centers and radii was the same as in MDITRE [23]. Briefly, centers and radii were initialized by performing K-means clustering (using KMeans from scikit-learn v1.3.2) for the set of detectors. Detector activation thresholds were initialized as the average over all subjects of the feature groups determined by KMeans. The prediction layer weights and biases were initialized from a standard Normal distribution. The rule and detector selectors were initialized to 0.5, as in [23]. As in our previous work [23], learning rates were set to be 0.001 for all parameters, except for taxa detector thresholds, ***η***^*T*^, which were set to be 0.0001. All temperature parameters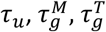, and *τ_q_*) were linearly annealed throughout training as described above. The model was trained for 5000 epochs for all datasets. Model convergence was defined as a decrease in training loss of less than within the last 100 epochs; in all cases convergence was achieved within 5000 epochs.

### Semi-synthetic data generation

Semi-synthetic data was generated from real data using a parametric bootstrapping procedure, similar to that in our prior work [22, 23] for generating longitudinal taxa counts, but modified to generate metabolites as well, and without temporal information. We simulated data from the 67 control samples of Wang et al. [4], which consist of metabolite levels measured with HPLC-MS/MS and taxa counts measured with shotgun metagenomic sequencing.

The overall procedure for generating semisynthetic data was as follows:

(1) Sample a set of subjects from the control subjects of Wang et al. and split the set into “cases” and “controls.”
(2) Sample a group (or groups) of metabolites and/or taxa to be perturbed.
(3) Calculate parameters for sampling the magnitude of synthetic perturbations, where the ranges of parameters were estimated from rules on real data.
(4) Generate data for “cases” and “controls” from the distributions determined in step 3.

Each of these steps is described in further detail below.

#### (1) Sampling sets of subjects

Subjects were uniformly sampled from the 67 control samples of Wang et al, with replacement if the number of subjects exceeded 67 in the simulation, and the set was then evenly split into “cases” and “controls.” Note that subjects were sampled in a nested manner to assure comparability of results across different sample sizes, i.e., 300 subject cases were sub-sampled from 1000 subject cases, 128 subject cases were sub-sampled from 300 subject cases, etc.

#### (2) Sampling groups of metabolites or taxa to be perturbed

The procedure for sampling metabolite groups was as follows:

1. Select a metabolite *p* uniformly at random from all the metabolites measured in the dataset, and obtain the metabolite *p*’s location in embedding space,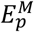.
2. Sample a radius *κ* from a Log-Normal distribution, with parameters set by the same procedure as described above for MMETHANE layer 1.
3. Calculate pairwise distances between all metabolites and 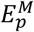, and select the group of metabolites to be perturbed as all those within distance *k* from 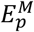.

To prevent the metabolite group from being unrealistically large, group sizes were restricted to a maximum of 15% of the total metabolites in the data; this was enforced via rejection sampling.

We followed the same procedure for sampling taxa groups to be perturbed as in our previous work [22, 23]. Briefly, clades were sampled uniformly from the tree with the restriction that clades contain a minimum of 5 members and a maximum of 30 members.

#### (3) Calculating perturbation magnitudes

##### Metabolites

To introduce a synthetic perturbation into the real metabolomics data, we sampled from a “control” distribution, 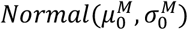, for a portion of subjects, and from a “case” distribution, 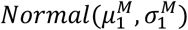, for the other subjects. We determined means of the distributions using the following procedure:

1. For each of the 6 datasets, determine the seed with the lowest loss for that dataset and obtain the set of MMETHANE rules for that seed. Over all the rules found in all the datasets, find the rule *k’* with the largest regression coefficient in absolute value.
2. Calculate the fold-changes for all detectors *j* in rule *k’*:

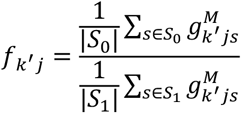 Here *S*_*0*_ is the set of control subjects and *S*_*1*_ the set of case subjects in the dataset that the rule was learned from, and 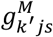 is the average of the log-transformed and standardized levels of metabolites in rule *k’*, detector *j*.
3. Choose the detector *j’* associated with the median fold-change, and calculate:

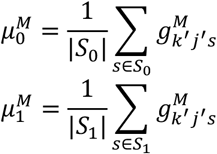

From this procedure, we found 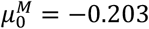 and 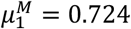. We set 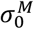 and 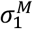 to 1.5, as in [22].

##### Taxa

We used the same values as in our previous work [22], 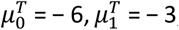, and 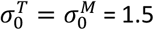.

#### (4) Generating data

##### Scenario 1: 1 metabolite group perturbed

For each metabolite *i* in the group to be perturbed, data was generated from a hierarchical model (top level modeling biological variation and the bottom level modeling measurement variability). Here, let *H* denote the set of metabolites selected to be perturbed in the chosen group and 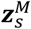 the standardized log-transformed values of all metabolites in subject *s*. To generate the semisynthetic data for each metabolite *m*, the following procedure was used:

1. Compute *p*_*sm*_:
  - if *m* is in *H* (the set of perturbed metabolites), sample:

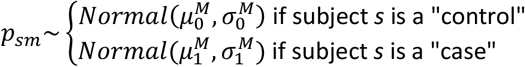
  - otherwise,

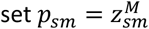
2. Sample measurement 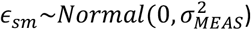. Here, 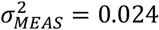 which we calculated from replicates from Dawkins et al [5].
3. Un-transform the data back to its original space:

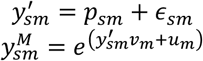

Here, *u*_*m*_ and *v*_*m*_ are the empirical mean and variance, respectively, of the untransformed levels of metabolite *m* over all subjects.

##### Scenario 2: 2 metabolite groups perturbed

This scenario is similar to scenario 1, except we separate the subjects in the “control” group into those receiving a perturbation in the first metabolite group (“control 1”), and those receiving a perturbation in the second metabolite group (“control 2”). Here, let *m*_*1*_ indicate the index of a metabolite in the first perturbation group *H*_*1*_, and *m*_*2*_ indicate the index of a metabolite in the second perturbation group *H*_*2*_. Then, we sample values for metabolite *m*_*1*_ as:

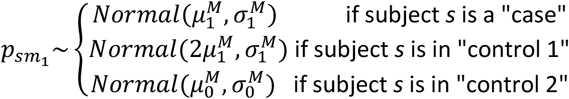

Similarly, for metabolite *m*_*2*_, we sample:

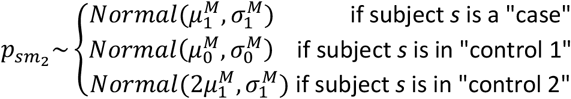

As in Scenario 1, if the metabolite is not in the group to be perturbed, *p*_*sm*_ is set to its original value in the data.

Note that these parameter settings (i.e., alternately doubling the magnitude of the perturbation for subjects in either of the control subgroups) ensure that the expected total amount of all perturbations is consistent in subjects.

##### Scenario 3: 1 taxonomic clade perturbed

We follow the same procedure as in our prior work [22]. For completeness, we describe it here as well. Let *C* denote the set of taxa selected to be perturbed in clade *c* and 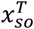 the relative abundance of taxa *o* for subject *s*.

1. For each subject *s*, sample a perturbation for the clade:

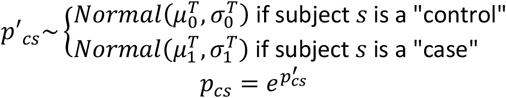
2. For each taxa compute *γ*_*so*_:
  - If taxa *o* is in *C, γ*_*so*_ is the proportion of the sampled perturbation corresponding to the relative abundance of taxa *o* in clade *c* in the original data:

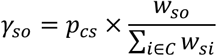
  - Otherwise, *γ*_*so*_ is set to the original relative abundance of taxa *o*:

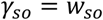
3. Add measurement noise in two stages:
  - First, sample from a truncated normal centered around *γ*_*so*_ with standard deviation set to 30% of the mean (*θ*=0.3)

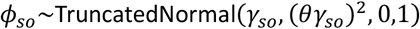
  - Second, generate counts from a Dirichlet-Multinomial distribution:

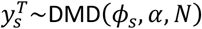

Here, *α* is set to 286 as in [22], and *N* is the total number of counts in subject *s*.

##### Scenario 4: 2 taxa clades perturbed

This scenario is analogous to Scenario 2, except using the taxa perturbation procedure described in Scenario 3.

##### Scenario 5: 1 taxa clade and 1 metabolite group perturbed

This scenario is analogous to Scenarios 2 and 4. We split controls into “control 1” and “control 2” groups. “Control 1” members then receive the taxa perturbation, “control 2” members receive the metabolite perturbation, and the “cases” receive both perturbations.

#### Data pre-processing and filtering

##### Processing metabolomics data

Metabolomics data was obtained from the original studies as feature intensities. These feature intensities were then filtered to remove metabolites in less than 15% of subjects, as in [5] and were then log-transformed and standardized.

##### Processing sequencing measurements

Sequencing measurements were re-processed from *fastq* files with the same bioinformatic tools to ensure consistency. For datasets with 16S rRNA amplicon sequencing, the raw data was reprocessed with DADA2 v1.20 [25] and the resulting amplicon sequencing variants were phylogenetically placed on a reference tree as described in [22]. For datasets with metagenomics sequencing, the raw sequences were processed with Metaphlan v3.0 [26], [56]. For both types of data, filtering was done as in [22]: taxa with counts below 10 in less than 10% of subjects were removed.

##### Processing datasets with multiple timepoints

Two datasets in our compendium, He et al. and Lloyd-Price et al., sampled multiple timepoints. However, the temporal information available in both datasets was insufficient to be useful for predictive modeling, as only the first two timepoints of He et al. showed any significant difference between groups and the time points sampled in Lloyd-Price et al. were substantially inconsistent across subjects. Thus, for both datasets, we selected one timepoint to use for analysis. For both He et al. and Lloyd Price et al., we used the timepoint that resulted in the largest sample size for each participant. For He et al., this was the timepoint at 2 months of life, and for Lloyd-Price et al., this was the first timepoint sampled from each participant.

#### Comparator machine learning methods

We benchmarked our model against lasso logistic regression (LR), random forest (RFs), adaptive boosting (AdaBoost), and feed forward neural network (FFNNs). For the ensemble benchmarking methods (RFs AdaBoost), data processing and filtering were performed identically as described MMETHANE. For LR and FFNNs, taxa relative abundances were transformed using the centered-log ratio transformation and then standardized. All methods used nested cross-validation, with the AUC as the metric, to tune hyperparameters. Nested 5-fold cross-validation was performed for all datasets, except the CDI dataset, on which nested leave-one-out cross-validation was performed due to the small sample size.

All comparator methods except FFNNs were implemented with Scikit-Learn (v1.3.2). LR parameter tuning was performed using LogisticRegressionCV, with the input settings and hyperparameter grid search performed as in [5]. RFs and AdaBoost parameter tuning was performed using GridSearchCV. For RFs, the hyperparameter grid to be searched over was set as in [5]. For AdaBoost, the hyperparameter grid was searched over 50 or 100 estimators and learning rates from 10^−2^ to 10^2^).

FFNNs were implemented in PyTorch, (v2.2.0) as a fully-connected, feed forward network with 3 layers. The number of nodes in each layer was scaled to the number of input features in the dataset according to the heuristics outlined in [63]. Models were run for 2000 epochs with a learning-rate of 0.001, dropout 0.1, and weight decay of 0.01. Models were assumed to have converged when the decrease in training loss over a window of 100 epochs was less than 0.01.

#### Determining modalities of predictors

For MMETHANE, predictor modality was directly determined from the model, using the rule set from the seed corresponding to the lowest total loss. For LR, we calculated the median odds and 95% interval of each feature over 10 seeds and 5 cross-validation folds within each seed, and retained every feature whose 95% odds-interval did not contain 0, as in [5]; modalities of retained features were then counted. We did the same for RFs and AdaBoost, but with Gini importance rather than odds. For FNNs, we calculated feature importances as the average integrated gradients using Captum (v0.7.0) [64], over ten seeds. Because no feature importances for the FNN were 0, we used the top 10 features for subsequent analyses.

### QUANTIFICATION AND STATISICAL ANALYSIS

The Mann Whitney U test from the Stats module of SciPy (v.1.11.3) was used to compare benchmarking results of methods on real and semi-synthetic data, with significance defined as *p*<0.05. All other details of statistical analyses and software can be found in the **Methods** section above, but are briefly summarized here. Calculation of the embedding dimension for taxa was performed using the KS test (p=0.05) from SciPy Stats (v.1.11.3). All machine learning models were run in Python (v3.11) using PyTorch (v2.2.0) and/or Scikit-Learn (v.1.3.2).

## Supplementary Figures and Tables

**Supplementary Figure 1.**
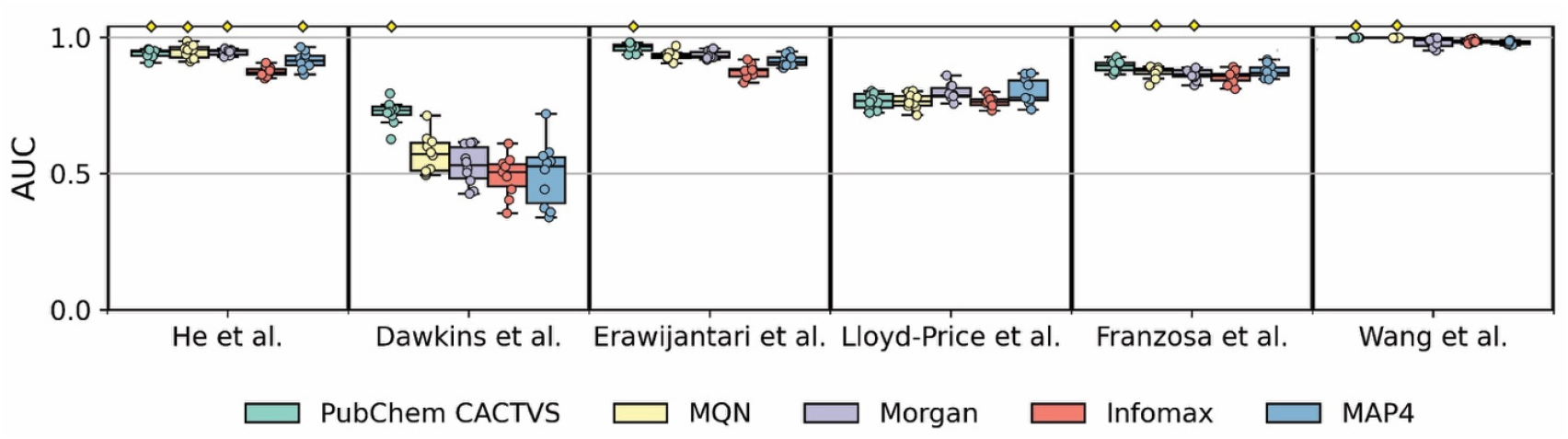
The PubChem CACTVS fingerprint yielded the most robust predictive performance when used as a molecular similarity measure for MMETHANE. Five-fold cross-validated AUC scores for prediction of host status on the six datasets in the compendium are shown, using five different molecular similarity measures (PubChem CACTVS, Morgan, MAP4, MQN and InfoMax). Box plots indicate medians and 95% intervals for runs over ten random seeds. Yellow diamonds indicate the top score or scores (if multiple scores were not significantly different from the top score, *p* > 0.05).

**Supplementary Table 1.**
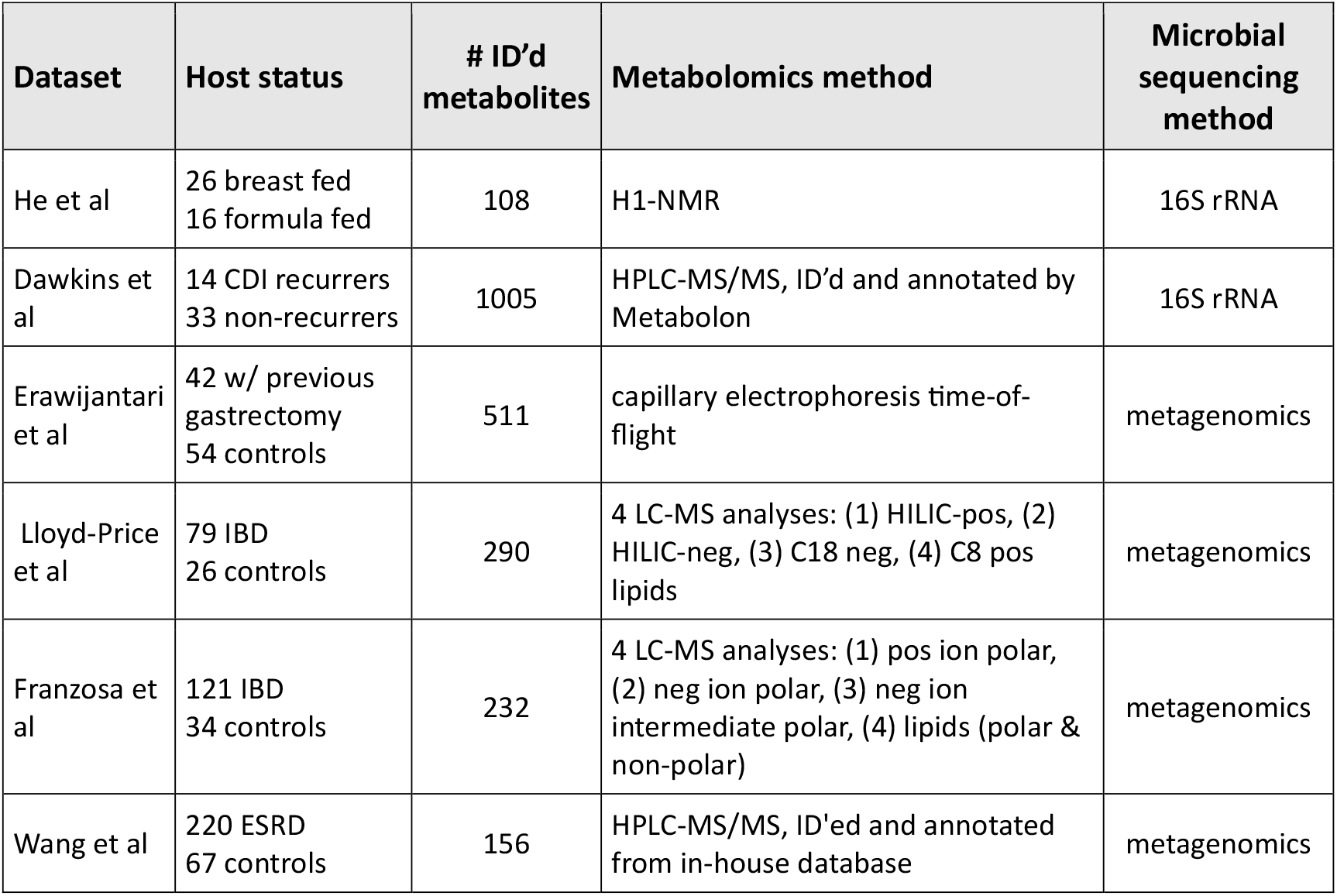
Compendium of paired microbial composition and metabolomics datasets. NMR = nuclear magnetic resonance. CDI = *Clostridioides difficile* infection. HPLC-MS/MS = high-performance liquid chromatography-tandem mass spectrometry. IBD = inflammatory bowel disease. LC-MS = liquid chromatography-mass spectrometry. ESRD = End-stage renal disease.

| Dataset | Host status | # ID'd metabolites | Metabolomics method | Microbial sequencing method |
| --- | --- | --- | --- | --- |
| He et al | 26 breast fed<br>16 formula fed | 108 | H1-NMR | 16S rRNA |
| Dawkins et al | 14 CDI recurrers<br>33 non-recurrers | 1005 | HPLC-MS/MS, ID'd and annotated by Metabolon | 16S rRNA |
| Erawijantari et al | 42 w/ previous gastrectomy<br>54 controls | 511 | capillary electrophoresis time-of-flight | metagenomics |
| Lloyd-Price et al | 79 IBD<br>26 controls | 290 | 4 LC-MS analyses: (1) HILIC-pos, (2) HILIC-neg, (3) C18 neg, (4) C8 pos lipids | metagenomics |
| Franzosa et al | 121 IBD<br>34 controls | 232 | 4 LC-MS analyses: (1) pos ion polar, (2) neg ion polar, (3) neg ion intermediate polar, (4) lipids (polar & non-polar) | metagenomics |
| Wang et al | 220 ESRD<br>67 controls | 156 | HPLC-MS/MS, ID'ed and annotated from in-house database | metagenomics |

**Supplementary Table 2.** Chemical fingerprints assessed and rationale for selection.

| Type | Fingerprint | # Dimensions | Numerical type | Reasoning |
| --- | --- | --- | --- | --- |
| Substructure keys | PubChem CACTVS [27] | 880 | Binary | Most comprehensive of substructure-keys |
|  | Molecular quantum numbers (MQN) [29] | 42 | Counts | Least complex of substructure-keys |
| Topological | Morgan (Extended connectivity fingerprint) [57] | 1024 | Binary | Commonly used as a comparison in other publications |
| Substructure + Topological | MAP4 (Substructure + Atom Pair) [59] | 1024 | Counts | Developed specifically for untargeted metabolomics |
| Deep learning | Infomax [30, 60] | 300 | Continuous $\in(0,1)$ | Highest performance in a comparative study |

**Supplementary Table 3.** Modality (metabolite or taxa) of predictors found when both metabolomics and taxa abundance data were given to models as inputs. LR = lasso logistic regression, RF = random forests, FFNN = feedforward neural networks. For LR, features were retained if their 95% cross-validated odds interval over 10 seeds did not contain 0. For RFs and AdaBoost, features were retained if their 95% cross-validated Gini importance over 10 seeds did not contain 0. For FNNs, the top 10 features were retained, ranked based on the mean of the integrated gradient of each feature over all subjects and 10 seeds.

|  |  | MMETHANE | LR | RF | AdaBoost | FFNN |
| --- | --- | --- | --- | --- | --- | --- |
| He et al | # Metabolite predictors | 1 | 3 | 2 | 1 | 10 |
|  | # Taxa predictors | 0 | 2 | 0 | 0 | 0 |
| Dawkins et al | # Metabolite predictors | 11 | 7 | 4 | 1 | 10 |
|  | # Taxa predictors | 0 | 0 | 0 | 0 | 0 |
| Erawijantari et al | # Metabolite predictors | 3 | 4 | 7 | 1 | 10 |
|  | # Taxa predictors | 16 | 1 | 2 | 0 | 0 |
| Lloyd-Price et al | # Metabolite predictors | 4 | 4 | 3 | 1 | 9 |
|  | # Taxa predictors | 21 | 0 | 0 | 0 | 1 |
| Franzosa et al | # Metabolite predictors | 2 | 4 | 9 | 2 | 10 |
|  | # Taxa predictors | 35 | 1 | 1 | 0 | 0 |
| Wang et al | # Metabolite predictors | 6 | 8 | 11 | 7 | 7 |
|  | # Taxa predictors | 18 | 4 | 2 | 0 | 3 |

